# Genetic identification of the RAS proteostatic machinery and its failure to regulate oncogenic variants

**DOI:** 10.64898/2025.12.10.693430

**Authors:** Johannes W. Bigenzahn, Felix Kartnig, Manuela Vollert, Vitaly Sedlyarov, Giulio Superti-Furga

## Abstract

The regulation of cellular homeostasis, differentiation, and proliferation is safeguarded by proteostatic mechanisms, which are crucial for maintaining cellular function. We used endogenously affinity-tagged KRAS cells in a fluorescence-activated cell sorting (FACS)-based CRISPR knockout screen to map genome-wide genetic requirements for proteostatic KRAS regulation. Three regulatory modules emerged, specifically cullin E3 ligase activity (*CUL3*, *NAE1*, *UBE2M*, *CAND1*, and *UBE2L3*), LZTR1 protein function (*LZTR1*, *NUDCD3*, and *ZRSR2*), and RAS GTPase modulation and processing (*NRAS*, *HRAS*, *FNTB*, *RCE1*, *ICMT*, and *GOLGA7*), all critical for regulating KRAS abundance. This expands our knowledge on the machinery controlling ubiquitin-meditated RAS family member regulation. Combining endogenous affinity tagging with genetic variant introduction, we found that the oncogenic KRAS G12D mutant exhibits reduced regulation by CRL3^LZTR1^, potentially contributing to oncogenic transformation and proliferation. In summary, our study genetically defines the machinery regulating RAS GTPase protein abundance, providing a foundation for a deeper molecular understanding and potential therapeutic exploitation of CRL3^LZTR1^-RAS GTPase regulation in human disease.

## Introduction

Proteostatic regulation of signaling proteins, thereby modulating their abundance and activity, is an important regulatory mechanism for maintaining proper cellular physiology including cellular survival, proliferation and differentiation(*1*, *2*). The RAS family of GTPase proteins encodes essential signaling hubs that transduce external and internal stimuli into effector responses. Specifically, the main group of RAS family GTPases, comprising KRAS, NRAS, and HRAS are well understood in their critical role in cancer cell initiation, maintenance, and proliferation(*3*). These proteins function as molecular switches, cycling between the GDP-bound “off” state and a GTP-bound “on” state to initiate downstream signal transduction. This process is further regulated by GTPase-activating proteins (GAPs), that promote the “off” state, and guanine nucleotide exchange proteins (GEFs), that facilitate the “on” state(*4*). Mutations in KRAS are among the most-frequently observed alterations in human cancers, particularly those affecting residues G12, G13 and Q61(*5*, *6*). These mutations result in a pool of mutant RAS GTPases predominantly locked in the GTP-bound “on” state.

Post-translational modifications of RAS proteins add crucial regulatory layers for precise signal control. These include the well-studied modifications in the C-terminal hypervariable region (HVR), such as sequential CAAX processing and palmitoylation, which are essential for intracellular trafficking and membrane localization, as well as various lysine-residue modifications, most notably ubiquitination(*7*).

Using various cellular models, we and others have identified the leucine zipper like post translational regulator 1 (LZTR1)-bound Cullin 3 (CUL3) E3 ubiquitin ligase complex (CRL3^LZTR1^) as a conserved proteostatic regulator of RAS GTPases(*8–11*). In this complex, LZTR1 acts as the substrate receptor, guiding the CUL3 E3 ligase scaffold to recruit the main RAS GTPases KRAS, NRAS and HRAS(*8*, *9*, *11*) as well as the closely related members MRAS(*10*, *11*) and RIT1(*10*) for targeted ubiquitination. Genetic inactivation of *LZTR1* can increase RAS GTPase protein abundance, resulting in enhanced RAS/MAPK pathway activation(*8–12*). Mutations in the *LZTR1* gene have been associated with a wide range of diseases and genetic disorders, including Noonan syndrome(*13*, *14*), a developmental disorder classified as a RASopathy(*15*), Schwannomatosis(*16*), and different cancers(*17*, *18*).

However, questions persist regarding the substrate specificity of LZTR1, particularly on its ability to bind and regulate the degradation and cellular localization of the main RAS GTPases, such as KRAS, in comparison to MRAS and RIT1, through ubiquitination-dependent mechanisms. To address these questions, we (1) employed protein-stability-reporter assays to validate the specificity and role of CRL3^LZTR1^ in regulating RAS GTPases abundance, (2) conducted a FACS-based genome-wide CRISPR screen to map the genetic requirements for endogenous KRAS abundance regulation by CRL3^LZTR1^, and (3) evaluated the impact of the frequently observed KRAS G12D and G12C oncogenic mutations on LZTR1-dependent proteostatic control in engineered isogenic cell lines. Collectively, these findings provide a more comprehensive understanding of CRL3^LZTR1^-based proteostatic regulation of RAS GTPases.

## Results

### The CRL3^LZTR1^ complex regulates the abundance of the main RAS GTPase KRAS as well as MRAS and RIT1

Various model systems have revealed distinct specificities of the CRL3^LZTR1^ complex toward different RAS GTPase family members. To clarify these differential abundance changes upon genetic inactivation of endogenous LZTR1 in an unbiased way, we performed a standardized, head-to-head comparison using a protein stability reporter (PSR) approach(*19*) (Fig. 1A). KRAS4A, as representative member of the main RAS GTPase family, along with MRAS and RIT1, were stably expressed in HAP1^EcoR^ and HEK293T^EcoR^ cells using a modified, bicistronic, protein abundance reporter vector with a dual HA tag (2HA) attached to the N-terminus of each GTPase protein (2HA-RAS GTPase-IRES-BlastR-P2A-GFP) (Fig. 1A). We opted for the smaller 2HA tag instead of the commonly used fluorescent protein tag to minimize potential artefactual changes in complex formation that could arise from the larger tag size. We also included a cDNA variant of KRAS4A (KRAS4A^(HD)^), which exhibits enhanced expression due to HRAS codon usage within the G-domain of KRAS4A(*20*). This variant had previously been used to characterize the cellular protein-protein interaction with endogenous LZTR1(*8*). Following PSR transduction, robust expression of all four GTPases was confirmed through flow cytometry and immunoblot analysis (Fig. 1B-E). Notably, while RIT1 exhibited uniform expression, both KRAS4A variants and MRAS displayed a biphasic HA staining distribution, indicating a subpopulation of cells with higher GTPase abundance. In contrast, the GFP reference control showed a homogenous expression pattern, confirming consistent reporter vector transduction (Fig. S1A-B). Upon genetic inactivation of LZTR1 using CRISPR/Cas9, we observed an increase in protein abundance for KRAS4A, MRAS, and RIT1 in both cell lines, confirming the ability of the CRL3^LZTR1^ complex in regulating not only MRAS and RIT1, but also the main RAS GTPase KRAS (Fig. 1B-E). Again, abundance of the GFP reference control remained unchanged, confirming the specificity of the observed effect (Fig. S1A-B). Interestingly, while we observed a homogenous increase in protein abundance for RIT1 by flow cytometry, both KRAS4A and MRAS proteins showed distinct patterns. In both cases, LZTR1 inactivation led to an increase in abundance within the lower-expressed subpopulation, while the higher-expressed fraction remained unchanged (Fig. 1D-E). Immunoblot analysis further confirmed a clear shift of RIT1 abundance upon LZTR1 inactivation, while changes of KRAS4A and MRAS were less pronounced (Fig. 1B-C). Interestingly, the highly expressed KRAS4A^(HD)^ variant showed no increase in abundance upon LZTR1 inactivation, suggesting potential saturation of the endogenous CRL3^LZTR1^ complex due to supraphysiological levels of exogenous RAS expression. In conclusion, these data demonstrate that CRL3^LZTR1^-dependent RAS abundance regulation is influenced by the substrate GTPase abundance, offering a possible explanation for previously reported differences in substrate specificity(*8–10*). Additionally, it exemplifies the value of flow cytometry-assisted protein abundance analysis, complementing immunoblot analysis, by providing crucial cellular (sub-) population resolution.

**Figure 1.**
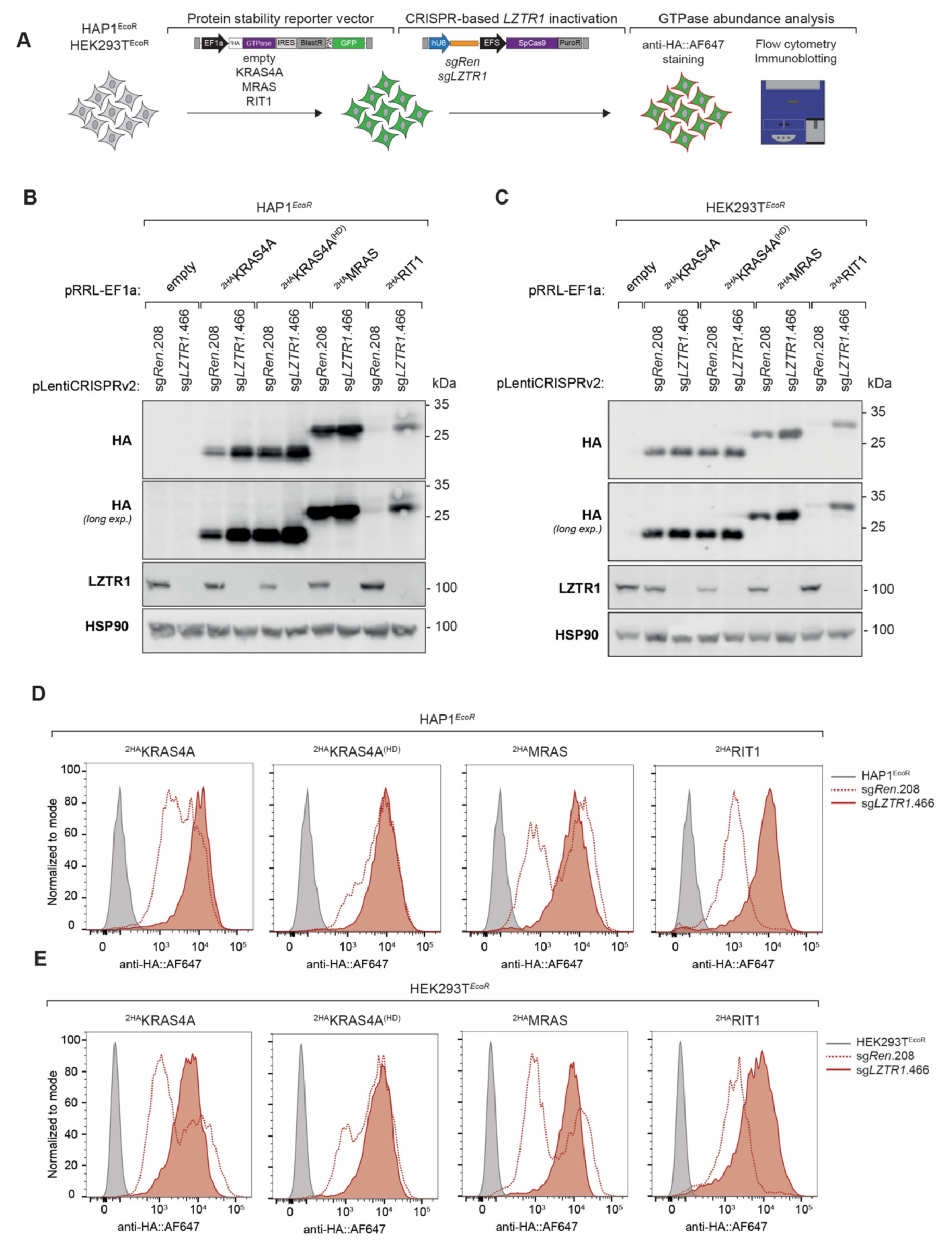
The E3 ligase complex CRL3^LZTR1^ regulates the abundance of different RAS family GTPases. **(A)** Schematic of protein stability reporter (PSR) vector-based evaluation of RAS family GTPase abundance alterations upon *LZTR1* inactivation. **(B-E)** Immunoblot and flow cytometric analysis with indicated antibodies of 2HA-tagged human RAS family GTPases KRAS4A (WT and HD variant), MRAS and RIT1 abundance upon CRISPR-based inactivation of *LZTR1* in HAP1*^EcoR^* **(B, D)** and HEK293T*^EcoR^* **(C, E)** cells. Immunoblot and flow cytometry results are representative of at least two independent biological experiments (n ≥ 2).

### Endogenous CRISPR tagging enables refined analysis of KRAS protein abundance

HAP1 cells exhibit robust expression of the CRL3^LZTR1^-subtrate GTPases KRAS, NRAS, HRAS, MRAS as well as RIT1 (Fig. S2A), and have been effectively utilized in the past to generate knock-in alleles with high efficiency(*8*, *21*). Given that overall KRAS expression levels impact CRL3^LZTR1^-dependent regulation, we generated HAP1 clonal cell lines harboring a 2HA tag at the N-terminus of endogenous KRAS, ensuring natural expression level and preservation of physiological regulation (Fig. 2A). Individual ^2HA^KRAS knock- in clones showed increased KRAS abundance following *LZTR1* knockout, as demonstrated by both immunoblotting and flow cytometry analysis (Fig. 2B-C displays results from LZTR1 knockout cell pools, while Fig. S2B shows individual *LZTR1* knockout clones), confirming the proteostatic regulatory role of LZTR1(*8*, *10*). Specificity of the endogenous tagging was confirmed through CRISPR-based inactivation of the tagged allele using a previously validated sgRNA targeting *KRAS* (Fig. 2B-C). Furthermore, reconstitution with *LZTR1* cDNA in individual ^2HA^KRAS *LZTR1*^KO^ clones resulted in decreased KRAS abundance, further corroborating the specific role of LZTR1 in proteostatic RAS GTPase regulation (Fig. S2C).

**Figure 2.**
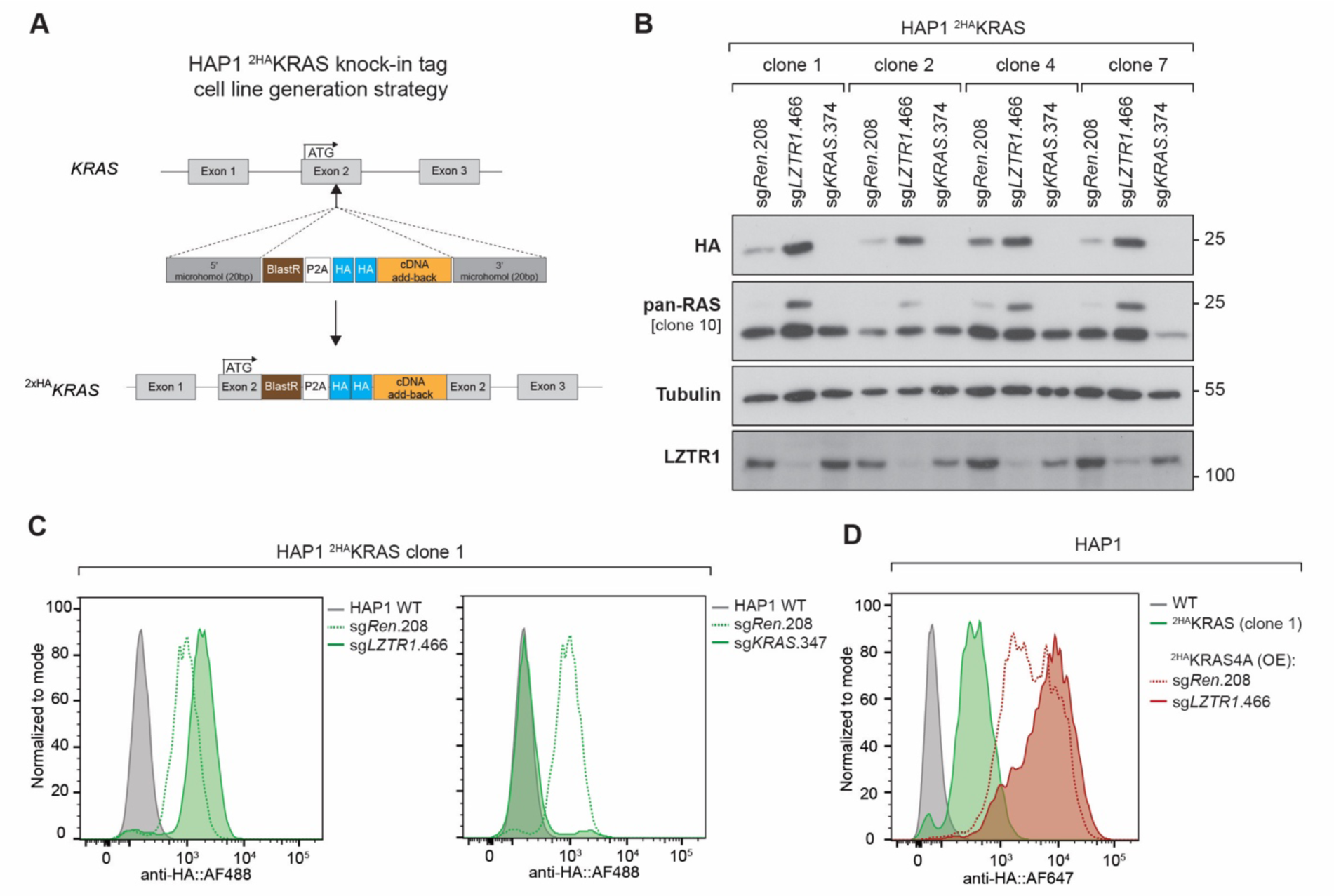
CRL3^LZTR1^ regulates the abundance of endogenous KRAS. **(A)** Schematic of microhomology-mediated end joining (MMEJ) CRISPR/Cas9-based 2HA tag knock-in cell line generation at the *KRAS* locus of HAP1 cells. **(B)** Immunoblot and **(C)** flow cytometric analysis of endogenous KRAS protein in HAP1 ^2HA^KRAS knock-in cell clones transduced with sg*Ren*.208, sg*LZTR1*.466 or sg*KRAS*.374. **(D)** Flow cytometric comparison of KRAS protein abundance in HAP1 ^2HA^KRAS knock-in cells and HAP1^EcoR^ ^2HA^KRAS4A overexpression cells transduced with sg*Ren*.208 or sg*LZTR1*.466. HAP1 WT cells serve as negative control. Immunoblot and flow cytometry results are representative of at least two independent biological experiments (n ≥ 2).

### FACS-based genome-wide CRISPR screen for cellular requirements of proteostatic KRAS regulation

Once established the role of LZTR1 in regulating RAS GTPases, we wanted to obtain an unbiased survey for any other genes that may act on the same or other proteostatic pathway. For this, we leveraged our endogenously tagged HAP1 reporter cell line model system to map cellular requirements for KRAS abundance regulation through FACS-based genome-wide CRISPR/Cas9 screening (Fig. 3A). Following sgRNA library transduction and expansion of the mutagenized cell pool, cell fractions displaying the lowest 5% (KRAS^LOW^) or highest 5% (KRAS^HIGH^) of KRAS protein levels, as determined by anti-HA tag immunostaining, were isolated by FACS. sgRNAs from the enriched cell fractions were subsequently quantified using sequencing (Fig. 3B and Fig. S3A-B). As expected, *KRAS* emerged as a top hit in the KRAS^LOW^ cell population, corroborating our previous specificity analysis in an unbiased manner and validating the robustness of our screening results (Fig. 3B).

**Figure 3.**
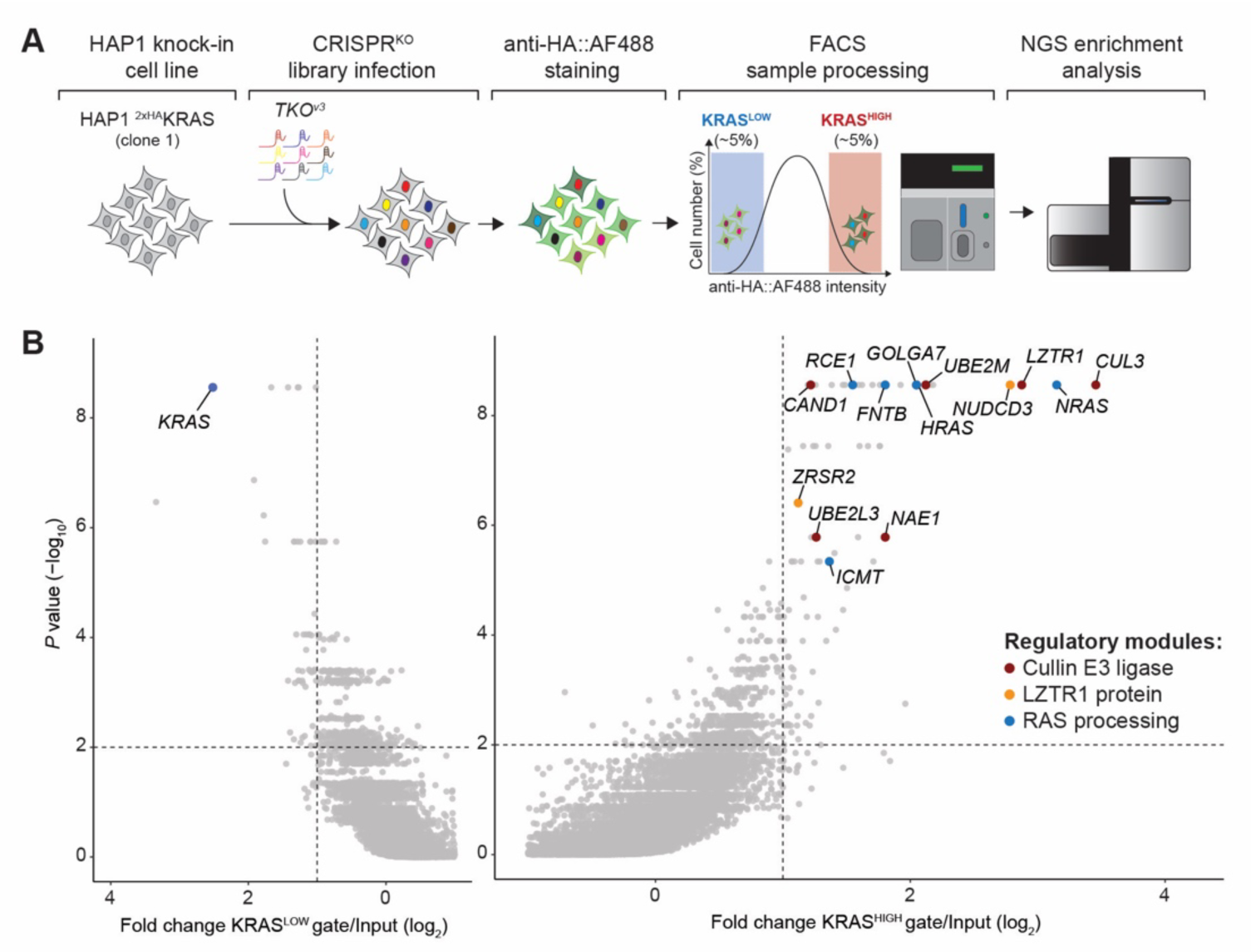
Functional genetic analysis of proteostatic KRAS regulation. **(A)** Schematic of FACS-based genome-wide CRISPR knockout screen in HAP1 ^2HA^KRAS knock-in cells. **(B)** Gene-level enrichment of sgRNAs identified in KRAS^HIGH^/Input and KRAS^LOW^/Input cell fractions representing negative and positive regulators of KRAS protein abundance respectively. Candidate genes with at least two-fold enrichment and a *P* value <0.01 as determined by MAGeCK RRA analysis were considered as significantly enriched. Genes involved in cullin E3 ligase, LZTR1 protein and RAS processing regulation are highlighted.

Shifting our focus to the top-ranking gene candidates in the KRAS^HIGH^ fraction with significant sgRNA enrichment, we identified 59 genes showing positive enrichment compared to unsorted cells (defined as a log_2_ fold change of >1 and a false discovery rate (FDR) <0.05) (Fig. 3B). Among the significantly enriched gene candidates in KRAS^HIGH^ cells (Fig. 3B and Fig. S4A), REACTOME pathway analysis revealed RAS processing, MAPK pathway regulation and E3 ubiquitin ligase-based protein ubiquitination as most significantly enriched processes (Fig. S4B). We identified *CUL3* and *LZTR1* as being strongly enriched, reaffirming their essential roles in the regulation of KRAS abundance. Several E3 ubiquitin ligases and associated proteins, including *BTRC* (bTrCP)(*22*), *RABGEF1* (Rabex-5)(*23*), *NEDD4*(*24*), and *WDR76*(*25*), have been previously reported in the literature as regulators of RAS protein abundance. Yet our genome-wide genetic screen in HAP1 cells only identified CRL3^LZTR1^ as prominent regulator of KRAS abundance albeit robust expression of all proposed RAS E3 ligases in HAP1 cells (Fig. S5A-B).

Furthermore, we were able to distinguish three distinct regulatory modules: cullin E3 ligase activity, LZTR1 protein function, as well as RAS GTPase protein modulation and processing (Fig. 3B and Fig. S3A-B). Enrichment of candidate-targeting sgRNAs was consistent when comparing KRAS^HIGH^ sorted fractions to both KRAS^LOW^ fractions and unsorted, mutagenized cells (Fig. S4A). All identified candidate genes displayed robust expression in HAP1 cells, as evidenced by mRNA expression levels, further supporting their phenotypic contribution (Fig. S6A). Gene essentiality can complicate the reliable identification of phenotypic modifier genes in genome-wide or targeted CRISPR screens. While several of our identified candidate genes exhibit general or context-specific essentiality across 1,150 cancer cell lines, none demonstrated marked cellular fitness defects at our selected screen time point (Fig. S3C-D). This suggests that our single time-point sorting strategy effectively assessed and captured essential candidate genes.

Within the cullin E3 ligase activity regulatory module, we identified the genes *NAE1* and *UBE2M*, both components of the E1 and E2 enzyme cascade responsible for conjugating the ubiquitin-like protein NEDD8 onto the cullin scaffold, thereby regulating its active state. Additionally, we identified *CAND1*, which functions as substrate receptor exchange factor. These findings provide strong genetic evidence supporting a direct ubiquitination-driven process for CRL3^LZTR1^-mediated RAS abundance regulation. Interestingly, we also identified one E2 ubiquitin-conjugating enzyme *UBE2L3*, which may act in concert with CRL3^LZTR1^. Comparative analysis of the identified candidate genes with a FACS-based CRISPR screen on proteostatic MYC oncogene regulation(*26*) demonstrated an overlap within the cullin regulatory module. LZTR1 protein function, as well as RAS GTPase protein modulation and processing, on the other hand, were specific to our KRAS screen (Fig. S6B). Based on existing literature we assigned *NUDCD3*, acting as Kelch-domain-specific HSP90 co-chaperone, and the splicing factor *ZRSR2*, which is involved in *LZTR1* mRNA processing, to the LZTR1 regulatory module. Interestingly, *ZRSR2* demonstrated weaker phenotypic enrichment compared to *NUDCD3* (Fig. 3B and Fig. S3A). Finally, the enrichment of sgRNAs targeting *FNTB*, *RCE1*, *ICMT* and *GOLGA7* highlighted the post-translational processing of the KRAS C-terminal HVR as a critical determinant for abundance regulation (Fig. 3B and Fig. S3B). Intriguingly, we also identified *NRAS* and *HRAS* expression as key determinants of CRL3^LZTR1^-mediated modulation of KRAS protein levels in our HAP1 cell line model (Fig. 3B and Fig. S3B). In summary, these data provide the first human genetic map of cellular regulators essential for adjustment of endogenous, natural KRAS abundance.

To confirm the robustness and reproducibility of our pooled, FACS-based screening results, we performed single-well validation experiments. HAP1 ^2HA^KRAS cells were infected with individual sgRNAs targeting the top-scoring negative regulators of KRAS abundance within the three identified regulatory modules. After sgRNA transduction, cells were immunostained, and KRAS abundance analyzed by flow cytometry (Fig. 4A). Genetic inactivation of candidate genes within the regulatory module governing CRL3^LZTR1^ E3 ligase function resulted in robust effects, which were concordant with our screening results (Fig. 4B). Within the RAS GTPase protein processing module, we observed marked increases of KRAS abundance upon knockout of the *KRAS* paralogues *NRAS* and *HRAS*, as well as the HVR-modifying enzymes *FNTB*, *RCE1*, and to a lesser extent *GOLGA7* (Fig. 4B). In contrast, *ICMT* inactivation, which showed a weak phenotypic enrichment in the screen (Fig. 3B and Fig. S3B), did not demonstrate a statistically significance increase (Fig. 4B). Taken together, our individual gene-focused validation results confirmed the robustness of our pooled screen findings and highlighted the central role of cullin E3 ligase and substrate receptor functionality, as well as RAS GTPase protein processing, in the proteostatic regulation of KRAS.

**Figure 4.**
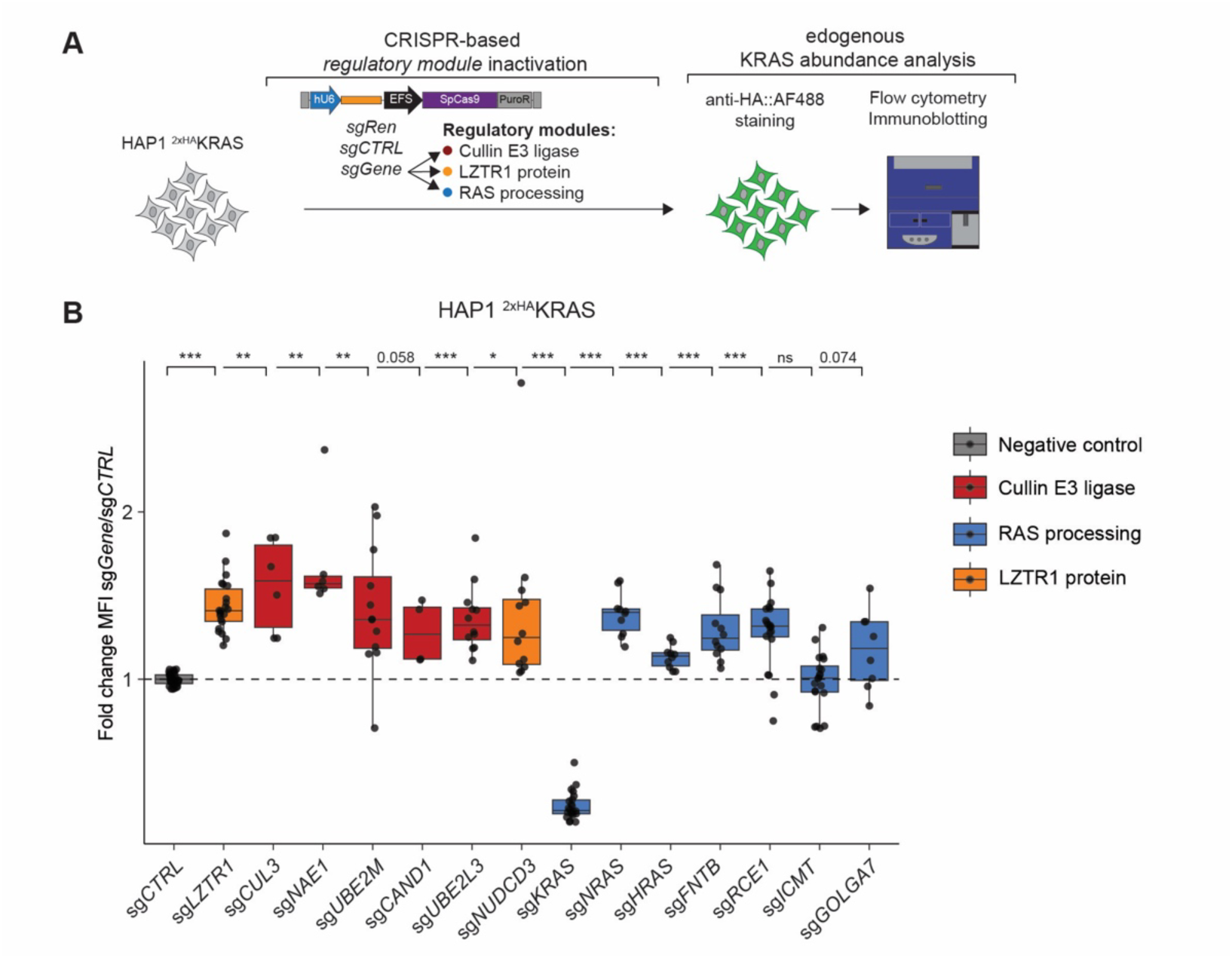
Distinct regulatory modules mediated CRL3^LZTR1^-based KRAS proteostatic regulation. **(A)** Schematic of flow cytometry-based validation of identified candidate genes. **(B)** Box and scatter plots displaying flow cytometric quantification of endogenous KRAS protein levels in HAP1 ^2HA^KRAS clone 1 cells transduced with negative control or sgRNAs targeting gene candidates within the individual regulatory modules as displayed in Fig. 3. Data are shown as fold change median fluorescence intensity (MFI) normalized to negative control sgRNAs. Each candidate gene is targeted by two independent sgRNAs and results are displayed by gene. Box plots show the median, lower and upper quartile with lower and upper whiskers extending to the furthest points within 1.5 times the IQR (inner quartile range). Each point represents an individual sgRNA and timepoint measurement. Data shown are obtained from at least two independent experiments (n ≥ 2). *, *P* < 0.05; **, *P* < 0.01; ***, *P* < 0.001; ns, not significant.

### KRAS G12D oncogenic mutant escapes CRL3^LZTR1^-based abundance regulation

Escape from CRL3^LZTR1^-based abundance regulation has been previously reported for mutations in the LZTR1 substrate GTPases MRAS and RIT1(*10*), that may contribute to their pathogenic effects in various cancers as well as Noonan syndrome. We therefore investigated whether a similar effect could also impact the proteostatic regulation of KRAS. We utilized our efficient knock-in system and introduced G12D or G12C mutations along with the 2HA tag into the endogenous KRAS locus (Fig. 5A). We selected these two mutations due to their high frequency in human cancers and their therapeutic importance(*27*). Our model provides the unique advantage of assessing the impact of individual mutations in an isogenic cellular setting at the endogenous expression level, while preserving both transcriptional and translational regulation. Interestingly, we observed increased KRAS abundance in cells harboring the G12D variant compared to WT cells, as shown by immunoblot as well as flow cytometry analysis. In contrast, cells with the G12C mutation displayed only a minor increase (Fig. 5B-C). Similarly, genetic inactivation of LZTR1 resulted in a pronounced increase in KRAS abundance in WT cells, whereas G12D mutant cells did not show any further increase. KRAS G12C expressing cells demonstrated an intermediate phenotype (Fig. 5D). These data provide evidence that certain KRAS mutations, such as G12D, exhibit reduced proteostatic regulation via CRL3^LZTR1^-mediated ubiquitination, potentially contributing to their oncogenic effect.

**Figure 5.**
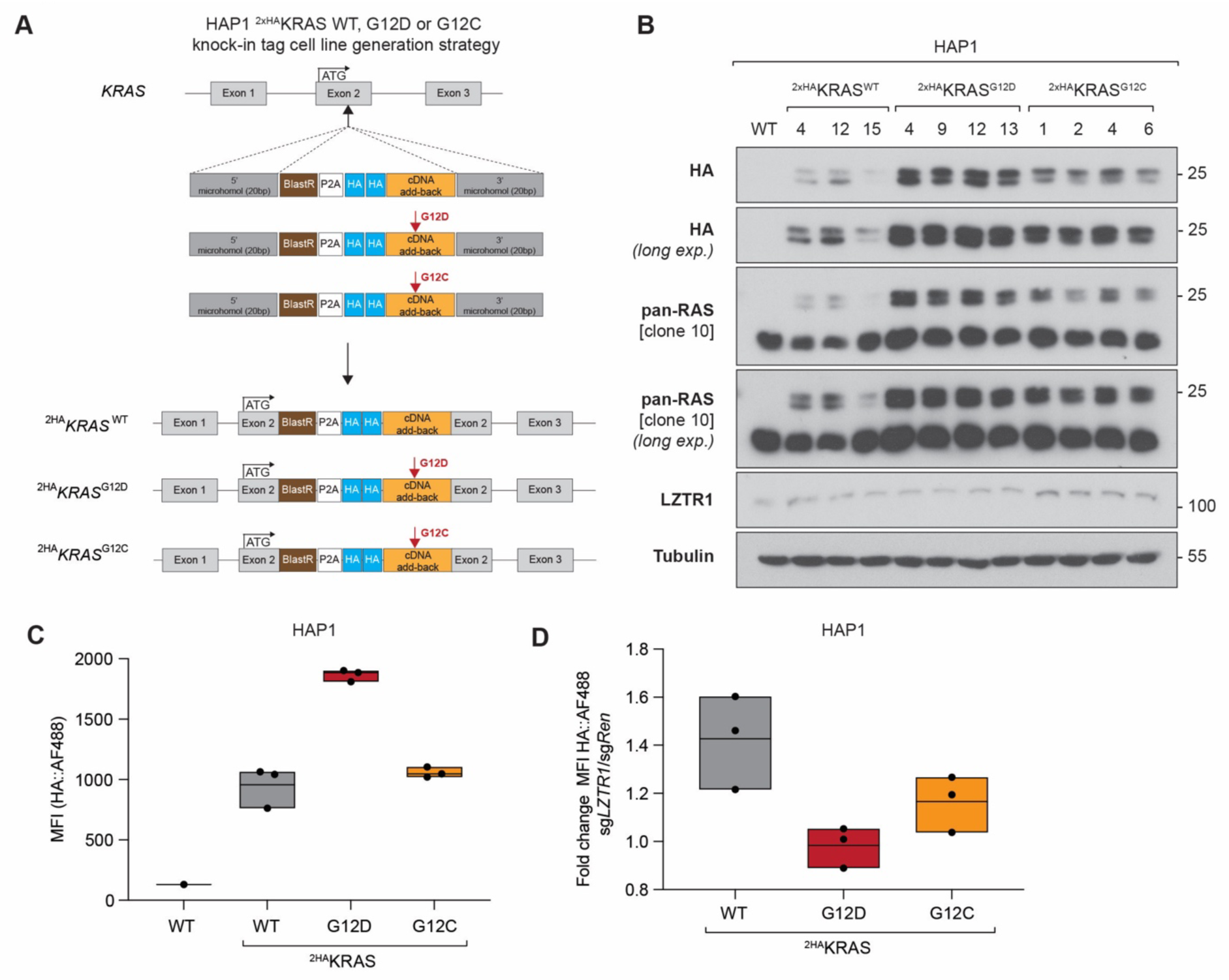
Oncogenic KRAS G12D escapes CRL3^LZTR1^-mediated protein abundance regulation. **(A)** Schematic of MMEJ CRISPR/Cas9-based 2HA knock-in tag and oncogenic mutation insertion at the endogenous *KRAS* locus of HAP1 cells. **(B)** Immunoblot and **(C)** flow cytometric analysis of endogenous KRAS protein in HAP1 ^2HA^KRAS knock-in WT (clone 4, 12, 15), G12D (clone 4, 9, 12) and G12C (clone 1, 2, 4) cell clones. HAP1 WT cells serve as negative control. **(D)** Flow cytometric analysis of endogenous KRAS protein in HAP1 ^2HA^KRAS knock-in WT (clone 4, 12, 15), G12D (clone 4, 9, 12) and G12C (clone 1, 2, 4) cell clones transduced with sg*Ren*.208 or sg*LZTR1*.466. Immunoblot and flow cytometry results are representative of at least two independent biological experiments (n ≥ 2). MFI, median fluorescence intensity.

## Discussion

The identification of the CRL3^LZTR1^ E3 ubiquitin ligase complex as a regulator of the RAS family GTPases K-, N-, and HRAS, along with MRAS and RIT1, provided evidence for an additional, conserved layer of RAS/MAPK signaling control, extending beyond the canonical modulation of the GTP/GDP “on”/”off” switch activity. This finding offered the first mechanistic explanation for the role of *LZTR1* mutations observed in various cancers and Noonan syndrome(*13*, *14*, *16*, *17*). However, uncertainty was raised regarding the specificity and GTPase substrate preference of LZTR1, as well as whether it actively drives their proteostatic regulation(*7*, *10*). In this study, we provide multiple lines of evidence demonstrating that CRL3^LZTR1^ regulates the abundance of both main as well as related RAS family GTPases and map the cellular genetic requirements for proteostatic KRAS regulation on a genome-wide scale. Furthermore, we show that the frequently observed KRAS G12D mutation escapes this abundance regulatory mechanism.

We addressed the question of abundance regulation using protein stability reporter-based monitoring to corroborate previous observations and clarify that both main RAS GTPases as well as MRAS and RIT1 depend on CRL3^LZTR1^ for their proteostatic regulation. Additionally, these experiments highlighted the important insight that basal GTPase expression levels are a critical determinant of LZTR1-based regulation, with strong overexpression potentially overriding or saturating the capacity of endogenous CRL3^LZTR1^ complexes. This finding might also explain the observed heterogeneity in the regulation of individual RAS GTPases across different cellular and tissue contexts(*28*, *29*). We therefore utilized our previously established experimental strategy of endogenous tagging of the KRAS gene to establish reporter HAP1 cell lines, which are amenable to FACS-based genome-wide CRISPR screening. This approach allowed us to further expand our molecular understand of the cellular requirements for CRL3^LZTR1^-based target regulation, by identifying several components within three distinct regulatory modules.

Identification of the cullin E3 ligase activity module mirrors functional genetic dependencies obtained by studying the abundance regulation of other human oncogenes, such as MYC(*26*), as well as the cellular sensitivity to molecular glue degraders(*30*, *31*) and protein targeting chimeras (PROTACS)(*31*). This underscores that CRL3^LZTR1^ exerts a ubiquitination-driven effect on RAS abundance. Beyond cullin complex activation and substrate receptor turnover, we further identified the E2 ubiquitin conjugating enzyme *UBE2L3* (UBCH7), that is known to collaborate with ARIH1 in ubiquitinating cullin ring ligase substrates, including CUL3-based complexes(*32–34*). Notably, *ARIH1* was not identified as a significant hit in our screen. Interestingly, *UBE2L3* has been identified as a fusion partner of KRAS in a prostate cancer cell line, displaying transforming activity in NIH3T3 cells and leading to endosomal localization of KRAS(*35*). However, *UBE2L3* resides in close genomic proximity to the LZTR1 locus (1Mb distance), raising the possibility of a CRISPR-mediated genome proximity-dependent collateral inactivation phenotype(*36*, *37*). Therefore, whether UBE2L3 represent a specific E2 directly collaborating with CRL3^LZTR1^ to regulate RAS ubiquitination will require further clarification.

Interestingly, in addition to the CRL3 complex itself, we identified *ZRSR2* and *NUDCD3* as specific limiting factors for proper KRAS abundance modulation. Previous research has shown that the minor spliceosome component ZRSR2 contributes to correct *LZTR1* mRNA splicing, thereby impacting protein expression(*38*). Consequently, mutational inactivation of *ZRSR2*, as observed in myelodysplastic syndromes (MDS), can lead to enhance RAS and RIT1 GTPase abundance due to reduced LZTR1 protein abundance, contributing to the transformation of hematopoietic cells. In contrast, NUDCD3 has been identified as a Kelch-domain containing protein-specific cochaperone for the HSP90 complex(*39*). Consistent with our findings, NUDCD3 acts as limiting factor for the CRL3^KBTB4^ complex in mediating the antiproliferative effect of the molecular glue degrader UM117(*40*). In this context, NUDCD3 was critically required for KBTBD4 protein stability. To our knowledge, our screen provides the second example of NUDCD3 being required for the function of Kelch-domain containing substrate receptor within the context of CUL3 E3 ligase-based ubiquitination. These findings highlight the importance of HSP90 cochaperone functionality in stability regulation and complex assembly. Moreover, the central importance of NUDCD3, potentially impacting many more Kelch-domain containing proteins, is illustrated by its broad essentiality for proliferation and survival in various cancer cell lines (Fig. S3C).

We further identified requirements beyond CRL3^LZTR1^ complex regulation on the side of RAS GTPase processing. As we have previously shown, modification of the C-terminal HVR tail of KRAS is important for the interaction with endogenous LZTR1(*8*). Here, our CRISPR screen provides orthogonal genetic evidence for these observations. The identification of the enzymes *FNTB*, *RCE1* and *ICMT*, responsible for generating the mature, membrane-associated form of RAS(*7*) genetically confirms the previous conjecture. Interestingly, we also identified *GOLGA7* as being required for KRAS abundance regulation. *GOLGA7* encodes a Golgi-located palmitoyltransferase subunit that has been shown to be specific for NRAS and HRAS, in contrast to KRAS(*41*). *GOLGA7* has also been identified as a context-specific essentiality in an NRAS G12D dependency screen(*42*). Therefore, the identification of *GOLGA7* as being required for CRL3^LZTR1^-based KRAS abundance regulation could be through its impact on the proper cellular localization of NRAS or HRAS.

We find the identification of *NRAS* and *HRAS* as genes affecting KRAS levels, highly interesting. This suggests that the expression levels of individual main RAS GTPases, in conjunction with potential RAS dimer formation or nano clustering(*43*) may impact CRL3^LZTR1^-based KRAS abundance regulation. The crosstalk between N- and HRAS isoform with KRAS and the impact on its oncogenic function has recently been described through *in vivo* CRISPR screening in a murine lung cancer model(*44*). Collectively, our genome-wide survey of proteostatic KRAS regulation has uncovered a multifaceted genetic interplay that is required for the proper regulation of KRAS abundance. This includes regulation of the cullin E3 ligase complex itself, as well as the stability, localization, and interaction capabilities of both the substrate receptor and target protein to elicit functional KRAS GTPase protein abundance regulation within the cell. It can be expected that the precise molecular interplay of these genetically defined components will uncover the regulatory relationships and how they are involved in cancer and RASopathies.

Finally, we addressed the question of whether a similar proteostatic regulatory mechanism governs the regulation of oncogenic KRAS mutants or if escape of proteostatic regulation may contribute to the oncogenic impact of RAS GTPases. Previously, MRAS and RIT1 have been identified as substrate GTPases, whereby mutations observed in Noonan syndrome would reduce or completely abrogate binding to LZTR1(*10*). This raised the question of whether a similar effect would be observable when looking at KRAS mutations. Preliminary evidence from the literature suggested that this might indeed be the case with respect to mutations such as G12D, G13D, and Q61H(*45*), as well as G12V(*11*, *46*). However, most of these data have been obtained under GTPase overexpression conditions and/or by using large affinity tags. Given the potential confounding factor of marked RAS overexpression on CRL3^LZTR1^-based regulation, we again employed our endogenous tagging platform to engineer two frequently observed oncogenic KRAS mutations: G12D and G12C. Interestingly, we observed a similar “escape” mechanism for the G12D mutant, while G12C behaved more like the WT protein. The discrepancy between G12D and G12C may be explained by differences in their intrinsic GTP hydrolysis rates(*47*) together with the overall low basal KRAS activation state present in the HAP1 cell line used(*48*). In line with this interpretation, mutation-specific phenotypic differences have similarly been observed in murine pancreatic organoid systems comparing distinct KRAS mutant alleles(*49*). Furthermore, very recent structural data indicate that LZTR1 binding of RAS GTPases is restricted to the inactive, GDP-loaded state(*50*), providing an additional mechanistic basis by which allele-dependent differences in nucleotide cycling could manifest as divergent phenotypes.

Our endogenous KRAS mutational data provides an experimental primer to study the vast array of different mutants in the context of LZTR1 regulation in human cells and raises the possibility that, similar to MRAS and RIT1, reduced binding of KRAS mutants to CRL3^LZTR1^ might contribute to their oncogenic effect. More broadly, it further emphasizes the critical requirement for keeping RAS GTPase protein levels under tight control, as abundance dysregulation can contribute to disease pathology(*5*).

In summary, the data presented in this manuscript using unbiased genome-wide genetic screening provide unconfutable evidence for CRL3^LZTR1^-mediated abundance regulation of both the main RAS GTPase KRAS as well as the closely related members MRAS and RIT1. This work presents the first genome-wide map of genetic requirements for CRL3^LZTR1^-based proteostatic RAS GTPase regulation. Our study offers a framework for a more precise molecular understanding of RAS abundance regulation, leveraging FACS-based genome-wide CRISPR screening as a powerful means to dissect the proteostatic regulation of signaling proteins. Undoubtedly, further mechanistic work will be necessary to fully untangle the specific cellular wiring that governs the abundance regulation of the proto-oncogene KRAS, mutated in around 25% of all human cancers. Overall, the deeper molecular understanding of proteostatic RAS GTPase regulation has the potential to lead to more refined mechanistic insights and may open new therapeutic avenues for RAS-driven human diseases.

## Materials and Methods

### Cell lines and reagents

HEK293T were obtained from ATCC (Manassas, VA, USA) and HAP1 cells were obtained from Horizon Genomics (Vienna, Austria). Cells were cultured in DMEM (D5796, Sigma-Aldrich, St. Louis, MO, USA) or IMDM medium (I6529, Sigma-Aldrich) supplemented with 10% (v/v) FBS (10437-028, Gibco, Grand Island, NY, USA, or S1810-500, Biowest, Riverside, MO, USA) and antibiotics (100 U/mL penicillin and 100 mg/mL streptomycin; P4333, Sigma-Aldrich). Cell lines were checked for mycoplasma infection by PCR or ELISA. Cell lines were authenticated by STR profiling.

### Antibodies

Antibodies used were: HA (#3724, Cell Signaling, Danvers, MA, USA), LZTR1 (E-12, sc-390166, Santa Cruz, Dallas, TX, USA), pan-RAS (clone 10, 05-516, Merck Millipore, Billerica, MA, USA), HSP90 (610418, BD Biosciences) and alpha tubulin (ab7291, Abcam). The secondary antibodies used were goat anti-mouse HRP (115-035-003, Jackson ImmunoResearch, West Grove, PA, USA), goat anti-rabbit HRP (111-035-003, Jackson ImmunoResearch), Alexa Fluor 488 goat anti-rabbit (A-11008, Thermo Fisher Scientific) and Alexa Fluor 647 goat anti-rabbit (A-21244, Thermo Fisher Scientific).

### Plasmids and cloning

For lentiviral expression of 2HA-tagged RAS GTPases, the cDNAs of human KRAS4A WT (NM_033360), KRAS4A^HD^ WT (KRAS-HRAS hybrid coding sequence design described in Lampson et al.(*20*) ), MRAS WT (isoform 1, NM_012219) and RIT1 WT (isoform 2, NM_006912) were PCR amplified from previously described pDONR221 entry plasmids using KOD DNA polymerase (71085, Merck Millipore) attaching a dual HA tag (2HA) at the N-terminus and inserted into the pRRL-EF1a- empty-IRES-BlastR-P2A-eGFP (LEIBG) proteostatic reporter vector using NEB DNA HiFi assembly (E5520S, NEB, Ipswich, MA, USA).

For endogenous tagging of the *KRAS* wildtype gene, a repair template containing BlastR-P2A-2HA flanked by 20bp micro homologies and PITCh sgRNA target sites was cloned by NEB HiFi DNA assembly (NEB) into the pCRIS MMEJ repair vector backbone (Addgene plasmid #63672) and an sgRNA targeting close to the KRAS start codon was cloned using GoldenGate assembly into a modified version of the pX458 vector (Addgene Plasmid #42230) containing an additional U6-PITCh sgRNA expression cassette. To generate endogenously tagged KRAS G12D and G12C oncogenic variants, the respective mutations were included into the cDNA add-back part of the repair template. Annotated sequences of the repair templates for PITCh KRAS knock-in v1 and v2 can be found in the corresponding supplementary sequence file. The sgRNA oligo sequences where as follows:

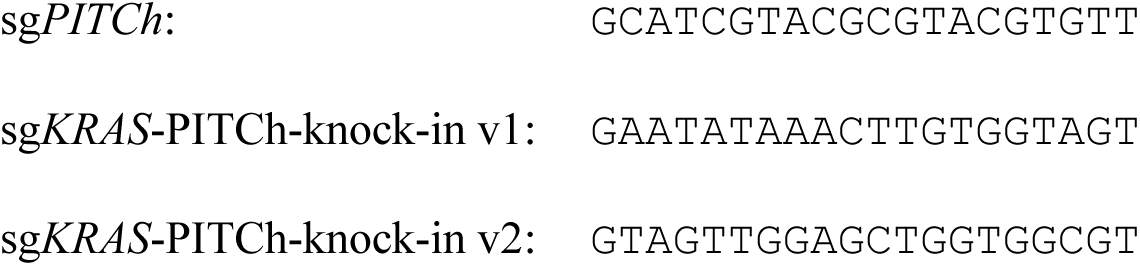

CRISPR/Cas9-based knockout cell line generation was performed using a derivative of the pLentiCRISPRv2 (Addgene plasmid #52961) lacking the SpCas9-attached FLAG epitope as described previously (Lc9v2nFP)(*8*). Individual sgRNAs were designed using CHOPCHOP(*51*) and CRISPick(*52*) or were selected from the Toronto knock-out library (TKOv3)(*53*). Oligonucleotides containing vector-compatible overhangs were annealed, phosphorylated using PNK (M0201S, NEB) and cloned into Lc9v2nFP using GoldenGate assembly with Esp3I (R0734S, NEB) and T4 DNA ligase (M0202S, NEB) followed by sequence verification using Sanger sequencing. sgRNAs targeting *Renilla luciferase* coding sequence (sg*Ren*.208: GGTATAATACACCGCGCTAC) or a save harbor locus on human chromosome 1 (sg*CTRL*-c1: GACTGAGGGTGGATAATCCA) were used as negative controls. Gene-targeting sgRNAs are labeled throughout the manuscript by gene name followed by the genomic targeting sequence position numbered according to the sequence position on the corresponding mRNA. sgRNA oligo sequences used in this study were as follows:

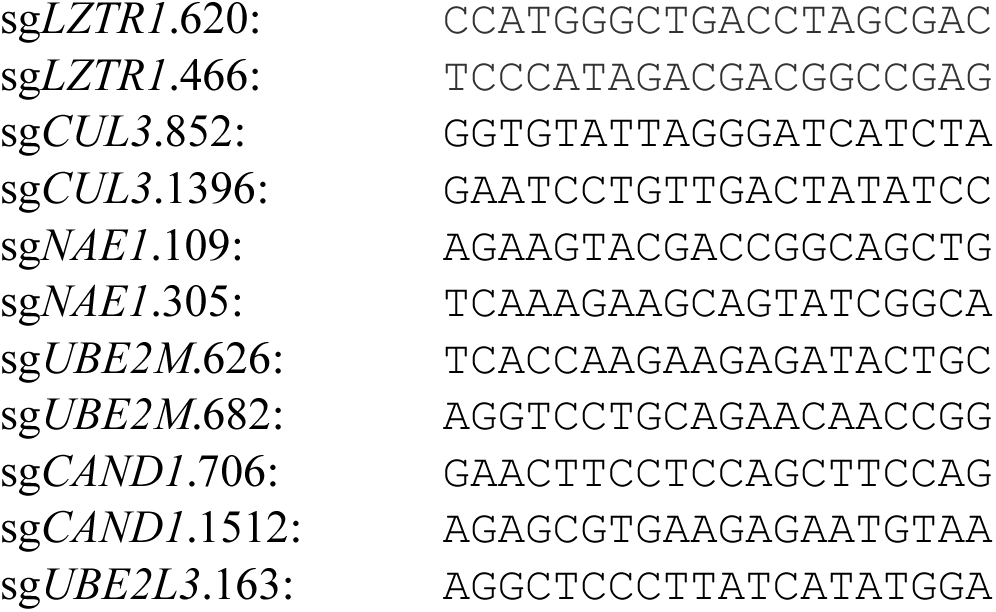
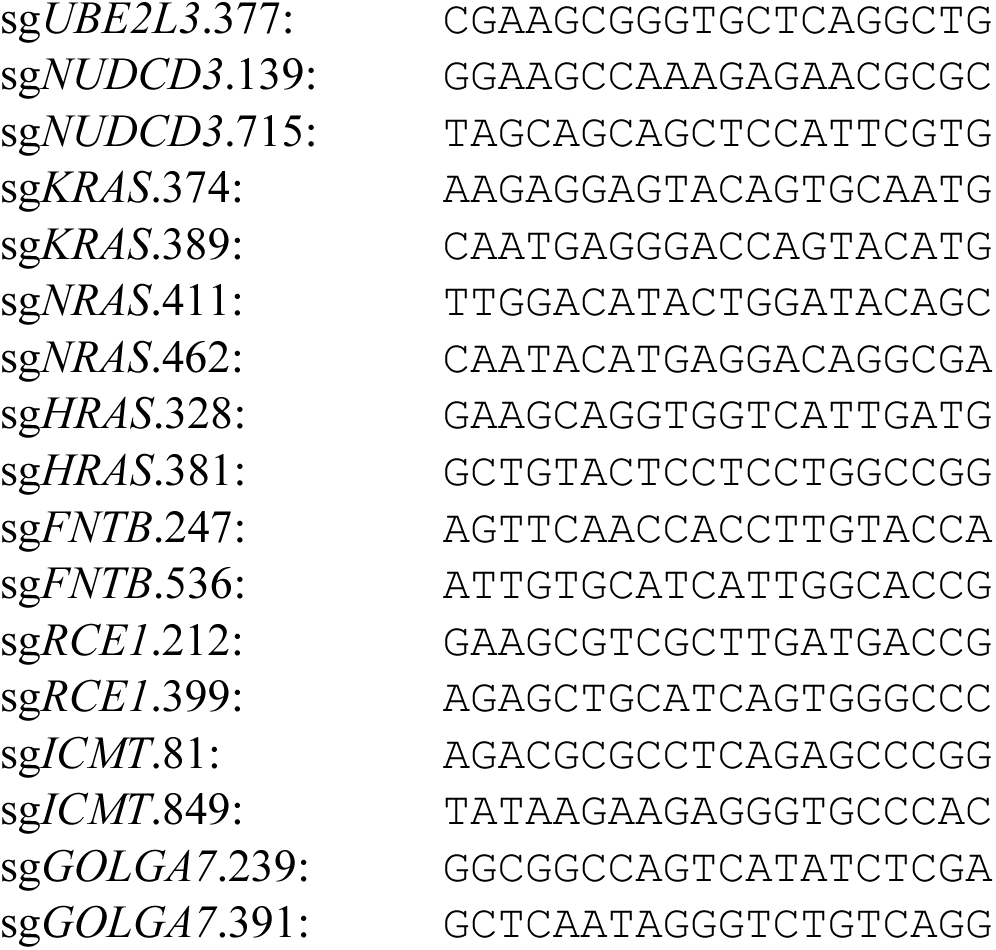

### Generation of CRISPR/Cas9-based knock-in cell lines

The generation of 2HA tag KRAS knock-in HAP1 cells was achieved by adapting a microhomology-mediated end joining (MMEJ)-mediated PTIChv2 CRISPR/Cas9-based targeting approach as described previously. HAP1 cells were transiently co-transfected with pX456-sgPITCh-sg*KRAS* PITCh-knock-in v1/v2 and pCRIS-PITChv2-BlastR-P2A-2HA-KRAS repair template vectors using TurboFectin 8.0 (TF81001, OriGene, Rockville, MD, USA). 3-4 days post transfection cells were treated with 20µg/mL blasticidin (ant-bl-05, InvivoGen, San Diego, CA) to enrich for clones that have successfully integrated the targeting constructs. Single cell clones were obtained by limiting dilution and verified by PCR using GoTaq2 DNA Polymerase (M7845, Promega, Madison, WI, USA) and Sanger sequencing as well as immunoblot analysis. Primer sequences to verify correct repair template insertion where as follows: sgKRAS_PITCh_knock_in_F: TGTAAAACGACGGCCAGACGATACACGTCTGCAGTCAAC sgKRAS_PITCh_knock_in_R: TCATGAAAATGGTCAGAGAAACC

M13 (sequence underlined) primer was used as sequencing primer.

### Lentiviral cell line generation

For lentiviral infections HEK293T cells were transiently transfected with psPAX2 (Addgene plasmid #12260), pMD2.G (Addgene plasmid #12259) and lentiviral expression vectors using polyethylenimine (PEI) transfection reagent (7854, BioTechne Tocris, Minneapolis, MN, USA). The medium was exchanged 6-12h after transfection and replaced with the respective target cell line specific medium. After 36-48h the virus-containing supernatant was harvested, filtered (0.45 µm), supplemented with 8µg/mL protamine sulfate (P3369, Sigma-Aldrich) and added to 40-60% confluent target cell lines. 24h after infection the medium was exchanged and replaced with fresh medium. Another 24h later, the medium was supplemented with the respective selection antibiotic to select for infected target cells. HAP1 and HEK293T cells were lentivirally transduced with pLERZIE (pRRL-EF1a-rtTA3-P2A-ZeoR- IRES-EcoR) to generate ecotropic receptor-expressing HAP1^EcoR^ and HEK293T^EcoR^ cells and selected with 200µg/mL zeocin (ant-zn-05, InvivoGen). For generation of proteostatic reporter vector expressing cells, pEcoEnv packaging plasmid was used instead of pMD2.G as described previously(*54*) and cells were selected with 20ug/mL blasticidin.

### Immunostaining for flow cytometry

To evaluate protein abundance changes of overexpressed RAS family GTPases or endogenous 2HA- tagged KRAS by flow cytometry, cells were harvested, washed twice with PBS (D8537, Sigma-Aldrich) and fixed with BD fix buffer I (557870, BD Biosciences, Franklin Lakes, NJ, USA) or 4% paraformaldehyde (PFA) for 20 minutes at room temperature (RT), washed twice with PBS containing 10% FCS (FACS buffer), followed by permeabilization using cold BD permeabilization buffer III (558050, BD Biosciences) or 90% MeOH for 30 minutes on ice. After washing twice in FACS buffer, cells were stained with anti-HA antibody (1:500-1:1000) for 1h or overnight at RT. Subsequently, cells were washed twice in FACS buffer and stained with goat anti-rabbit Alexa Fluor 488 (AF488, 1:1000) or goat anti-rabbit Alexa Fluor 647 (AF647, 1:1000) for 1h at RT, and subsequently cells were washed three times and resuspended in FACS buffer for flow cytometric analysis on an LSR Fortessa (BD Biosciences). Data analysis was performed using FlowJo software version v7 and v10 (BD Biosciences). The gating strategy for flow cytometry analysis is shown in Fig. S5.

### Immunoblotting

Cells were lysed using Nonidet-40 lysis buffer (50mM Tris-HCl pH 7.5, 150mM NaCl, 5mM EDTA, 1% NP-40, and one tablet of Roche EDTA-free protease inhibitor cocktail (11873580001, Sigma-Aldrich) per 50 mL) for 10 min on ice. Lysates were cleared by centrifugation (13000 rpm, 10 min, 4°C). The proteins were quantified and normalized with Bradford assay (#5000006, Bio-Rad, Hercules, CA, USA) using γ-globin as a standard (#5000005, Bio-Rad). Cell lysates were resolved by SDS-PAGE and transferred to nitrocellulose membranes Amersham^TM^ Protran^TM^ 0.45 µm (10600002, GE Healthcare, Little Chalfont, UK). The membranes were immunoblotted with indicated antibodies and bound antibodies were visualized with horseradish peroxidase–conjugated secondary antibodies using the Pierce^TM^ ECL western blotting substrate (32106, Thermo Fisher Scientific) or Clarity^TM^ Western ECL substrate (#170-5060, Bio-Rad) and using Amersham^TM^ Hyperfilm™ ECL™ films (28906837, Marlborough, MA, USA) or ChemiDoc^TM^ MP imaging system (BioRad).

### Fluorescence activated cell sorting (FACS)-based CRISPR-Cas9 screen

The pooled genome-wide CRISPR screen was performed by lentivirally transducing HAP1 ^2HA^KRAS clone 1 cells with the Toronto KnockOut CRISPR Library - Version 3 (TKOv3)(*53*) at low multiplicity of infection (MOI) and 500- to 1,000-fold library representation. Library-transduced cells were selected with puromycin (1μg/mL, ant-pr-1, InvivoGen) and expanded for eight days to allow for successful editing while maintaining 1,000-fold library representation. At time of cell harvest for immunostaining and subsequent FACS sorting, unsorted and unstained control samples were harvested corresponding to 1,000-fold library representation, snap-frozen and stored at -80°C until library processing. For the sorting-based screen, cells were harvested, washed twice with PBS and fixed with BD fix buffer I for 20 minutes at room temperature (RT), washed twice with PBS containing 10% FCS (FACS buffer), followed by permeabilization using cold BD permeabilization buffer III for 30 minutes on ice. After washing twice in FACS buffer, cells were stained with anti-HA antibody (1:500) for 1h at RT, washed twice in FACS buffer, stained with goat anti-rabbit Alexa Fluor 488 (AF488, 1:1000) for 1h at RT, and thereafter cells were washed three times and resuspended in FACS buffer. Before sorting cells were strained through a 40 μm mesh and the 5% of cells with the lowest and the highest anti-HA::AF488 signal, corresponding to endogenous KRAS protein abundance, were sorted into FACS buffer using a Sony SH800 cell sorter (Sony, Minato, Japan). The gating strategy for flow cytometric cell sorting is shown in Fig. S5. Sorted cell pellets were snap-frozen and similarly stored at -80°C until library processing.

### Generation of next-generation sequencing libraries

Genomic DNA was extracted using the DNeasy Blood & Tissue Kit (69506, Qiagen, Ipsogen SA, USA). Fixed and sorted samples were additionally de-crosslinked in PBS and lysis buffer AL (Qiagen) with the addition of proteinase K overnight at 56 °C with agitation. Subsequently the sgRNA cassette in all samples was amplified using a two PCR protocol based on Hart et al.(*53*) and Sanson et al.(*52*). The first PCR was performed using ExTaq DNA polymerase (RR001B, TaKaRa, Kyoto, Japan) in multiple 50μL reactions and 5μg of genomic DNA (gDNA) as input per PCR reaction for unsorted control cell populations and 1μg gDNA for sorted cell populations using the following cycler conditions: 1min at 95°C, 24 (5μg input samples) or 25 (1μg input samples) cycles of 30sec at 95°C, 30sec at 58°C, and 40sec at 72°C followed by a final extension of 10min at 72°C. The resulting PCR products were purified using AmpliClean™ magnetic beads (AP-005, NimaGen, Nijmegen, NL) and 20ng purified PCR product were used as template for a second, barcoding PCR, attaching standard Illumina adapters using the following cycler conditions: 1min at 95°C, 10 cycles of 30sec at 95°C, 30sec at 58°C, and 30sec at 72°C followed by a final extension of 10min at 72°C. The resulting libraries were pooled, purified using AmpliClean™ magnetic beads sequenced on an Illumina HiSeq system at the Biomedical Sequencing Facility (https://biomedical-sequencing.at/). Primers used for library preparation are listed in Table S1.

### Analysis of pooled CRISPR screens

Sequences of sgRNAs were extracted from NGS reads, aligned to the TKOv3 sgRNA library reference file, counted and normalized using an in-house Python script(*55*). Subsequently, normalized count tables were used as input for MAGeCK (0.5.9)(*56*) to calculate average log^2^ fold changes (LFCs), *P* values and false discovery rates (FDRs). MAGeCK-based CRISPR screen results of KRAS abundance regulation are provided in Table S2.

## Data analysis

Experimental sample number and the number of biological replicates is provided in the figure legends. Data organization and calculations were performed using Microsoft Excel (Microsoft, Redmond, WA, USA) or R programming environment within RStudio (Posit PBC, Boston, MA, USA). Data plotting was done using GraphPad Prism 9 (GraphPad Software, Boston, MA, USA) or RStudio. Bioinformatic analysis of the CRISPR screen is described above. The source of published data sets for comparative mRNA expression and CRISPR screen analysis is provided in the corresponding figure legends. Gene effect CHRONOS score are derived from the DepMap consortium portal based on 24Q2 data release(*57*, *58*). Statistical significance of flow cytometry-based protein abundance analysis was calculated with a pairwise t-test with Bonferroni-based multiple comparisons test correction. REACTOME-based pathway enrichment analysis was performed using ENRICHR(*59*) and results are provided in Table S2.

## Data availability

All data needed to evaluate the conclusions in the paper are present in the paper and/or the Supplementary Materials. CRISPR screen NGS raw data will be made publicly available upon publication through the Sequence Read Archive (SRA) database.

## Acknowledgment

We thank all members of the G.S.-F. laboratory, L. Heinz, G. Winter, C. Mayor-Ruiz, M. Jäger and B. Mair for discussion and feedback; U. Goldmann for help with data handling and storage; the Biomedical Sequencing Facility (CeMM/Medical University of Vienna) for the NGS sequencing. Plasmids and CRISPR library obtained through Addgene were a gift from D. Trono, F. Zhang, J. Moffat and T. Yamamoto.

## Author contributions

J.W.B. and G.S.-F. designed research; J.W.B, F.K. and M.V. performed research; V.S. performed bioinformatic analysis; J.W.B. and G.S.-F. analyzed and interpreted the data; J.W.B. and G.S.-F. wrote the paper with contributions from all other co-authors.

## Table of contents for supplementary materials

Figs. S1 to S7 Tables S1 to S2

## Supplementary information

