## Supplementary information for "Genetic identification of the RAS proteostatic machinery and its failure to regulate oncogenic variants"

### Supplementary materials:

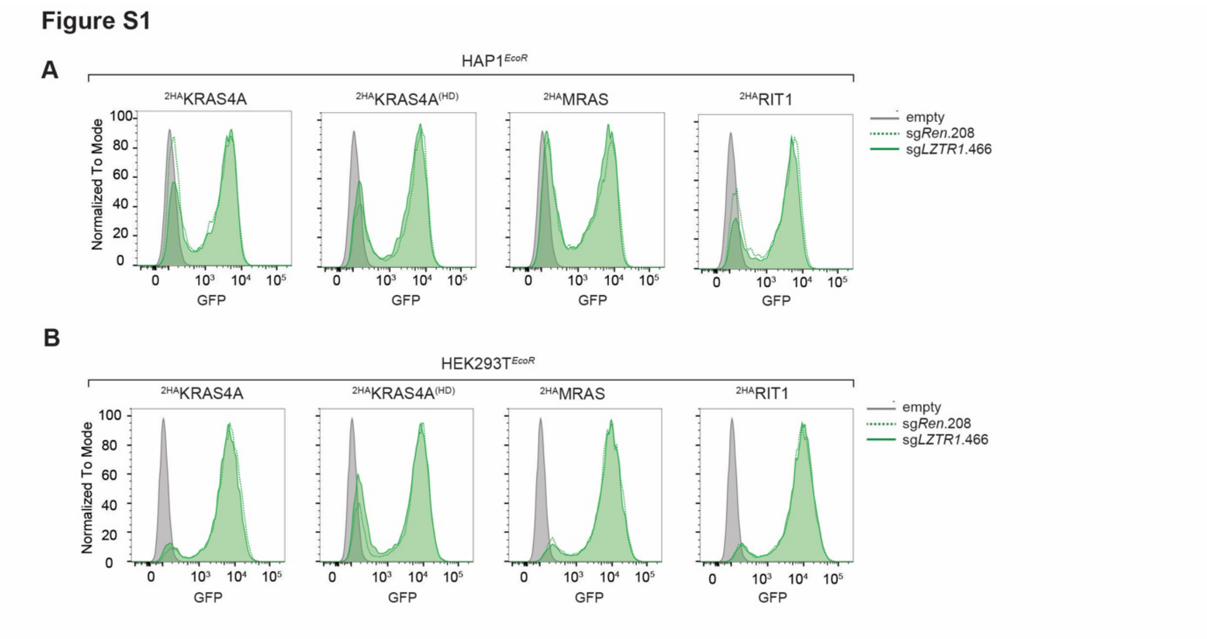


**Figure S1. The E3 ligase complex CRL3$^{LZTR1}$ regulates the abundance of different RAS family GTPases. (A-B)** Flow cytometric analysis of protein stability reporter (PSR) vector-transduced HAP1$^{EcoR}$ **(A)** and HEK293T$^{EcoR}$ **(B)** cells expressing different RAS family GTPase proteins transduced with sg*Ren*.208 or sg*LZTR1*.466. Flow cytometry results are representative of at least two independent biological experiments (n ≥ 2).

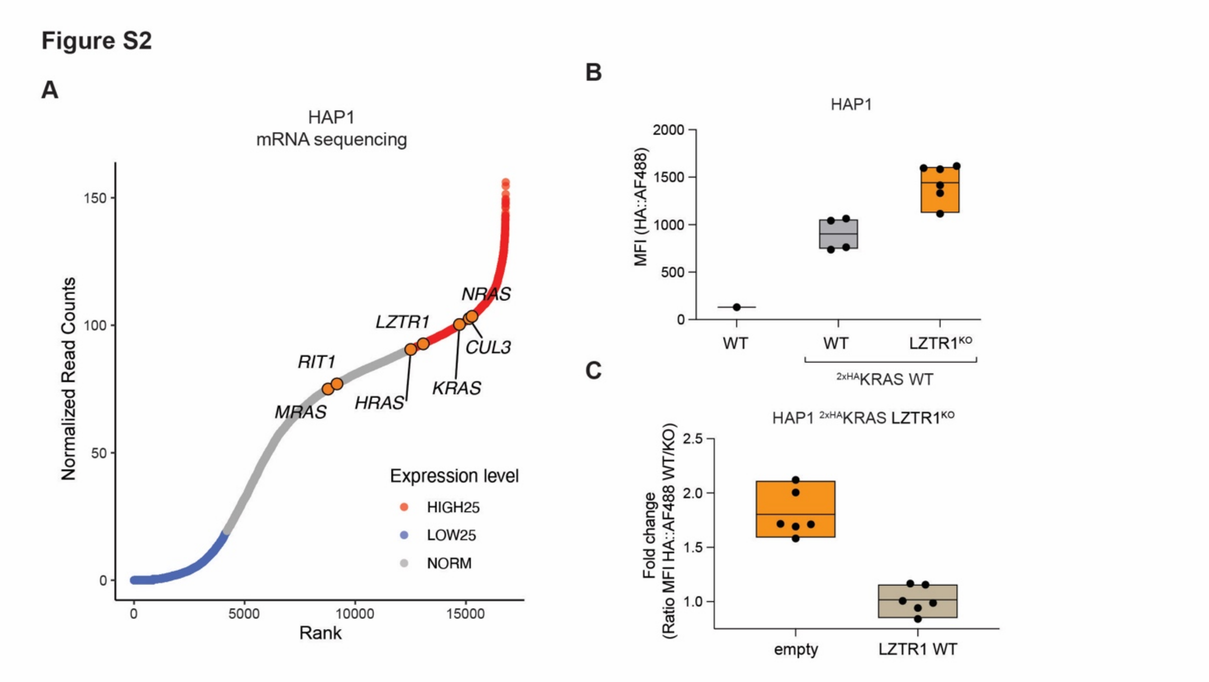


**Figure S2. CRL3$^{LZTR1}$ regulates the abundance of endogenous KRAS. (A)** Ranked expression of CRL3$^{LZTR1}$ and corresponding substrate RAS GTPase genes in HAP1 cells. mRNA expression is derived from published RNA sequencing data*(48)*. **(B)** Flow cytometric quantification of endogenous 2HA-tagged KRAS protein levels in control or LZTR1 knockout single cell clones. HAP1 WT cells serve as negative control. **(C)** Flow cytometric quantification of 2HA-tagged KRAS protein levels in LZTR1 knockout clones after lentiviral transduction with empty vector or *LZTR1* WT cDNA. Data in **(B)** and **(C)** are representative results of at least two independent biological experiments ($n \geq 2$). MFI, median fluorescence intensity.

### Figure S3

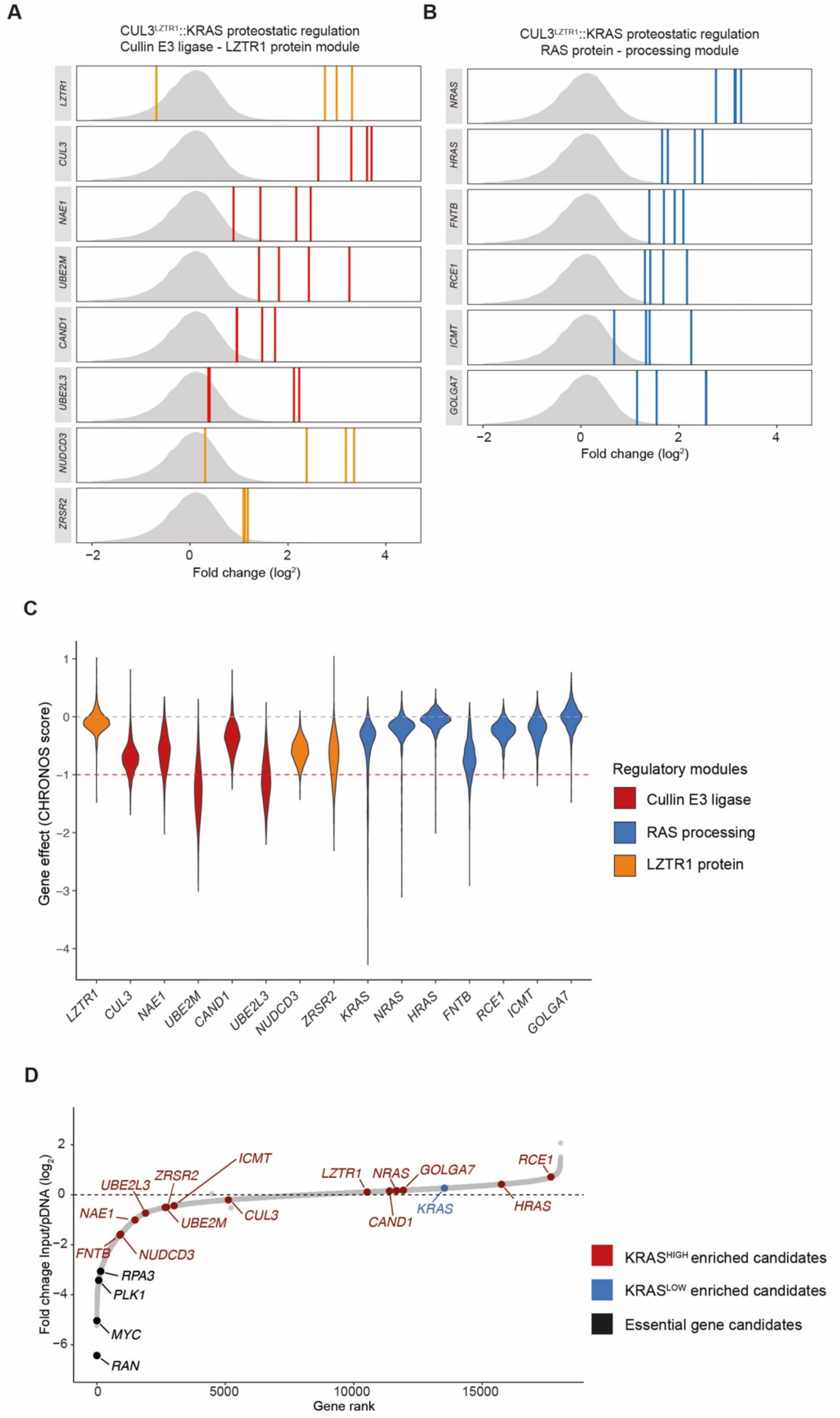
A
CUL3LZTR1::KRAS proteostatic regulation
Cullin E3 ligase - LZTR1 protein module
LZTR1
CUL3
NAE1
UBE2M
CAND1
UBE2L3
NUDCD3
ZRSR2
−2
0
2
4
Fold change (log2)
B
CUL3LZTR1::KRAS proteostatic regulation
RAS protein - processing module
NRAS
HRAS
FNTB
RCE1
ICMT
GOLGA7
−2
0
2
4
Fold change (log2)
C
Gene effect (CHRONOS score)
1
0
−1
−2
−3
−4
LZTR1
CUL3
NAE1
UBE2M
CAND1
UBE2L3
NUDCD3
ZRSR2
KRAS
NRAS
HRAS
FNTB
RCE1
ICMT
GOLGA7
Regulatory modules
Cullin E3 ligase
RAS processing
LZTR1 protein
D
Fold chnage Input/pDNA (log2)
2
0
−2
−4
−6
ICMT
UBE2L3
ZRSR2
NAE1
UBE2M
CUL3
FNTB
NUDCD3
RPA3
PLK1
MYC
RAN
LZTR1
NRAS
GOLGA7
CAND1
KRAS
RCE1
HRAS
0
5000
10000
15000
Gene rank
KRASHIGH enriched candidates
KRASLOW enriched candidates
Essential gene candidates

**Figure S3. sgRNA enrichment and gene essentiality of candidate genes mediating proteostatic KRAS regulation (A-B)** Fold change of individual sgRNAs for genes of interest significantly enriched in the KRAS$^{HIGH}$ fraction. Bars represent the mean $log_2$ fold change of each sgRNA, and density plots display the overall distribution of all sgRNAs in the KRAS$^{HIGH}$ fraction compared to unsorted control. sgRNAs are colored based on their grouping into the regulatory modules cullin E3 ligase, LZTR1 protein and RAS processing regulation. **(C)** Distribution of CRISPR knockout-based gene inactivation effects (CHRONOS score) across 1150 cancer cell lines derived the from Broad Institute DepMap Consortium (24Q2, public)(*57*). Gene candidates are colored based on their grouping into the regulatory modules cullin E3 ligase, LZTR1 protein and RAS processing regulation. **(D)** Gene-level sgRNA abundance changes in HAP1 $^{2HA}$KRAS clone 1 cells comparing unsorted cell populations harvested at the screen time point compared to sgRNA abundance in the library plasmid pool. Selected candidates are labeled based on their respective enrichment properties and common essential gene candidates are depicted as positive depletion control reference.

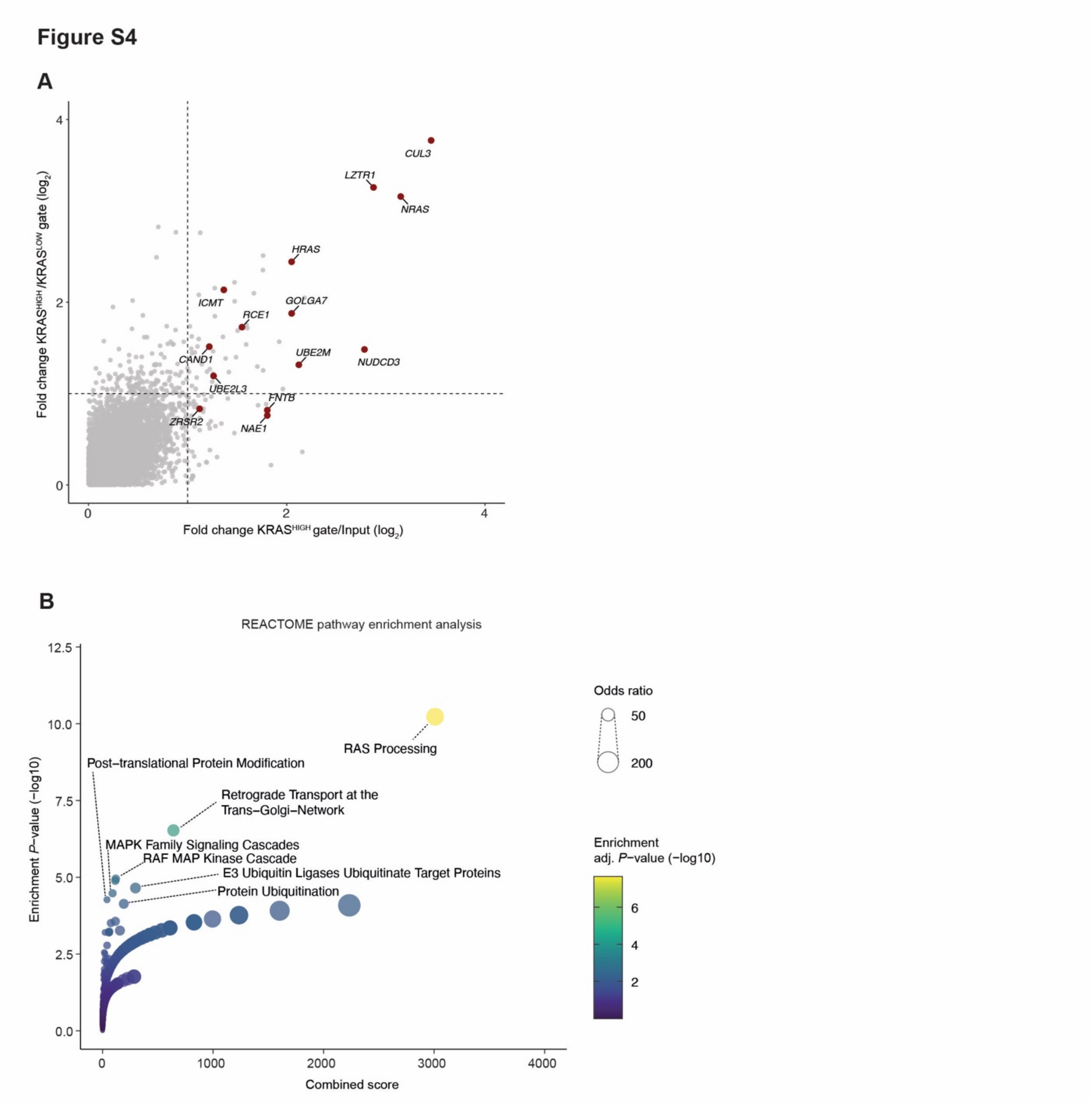


**Figure S4. Functional genetic analysis of proteostatic KRAS regulation. (A)** Gene-level enrichment of sgRNAs identified in KRAS$^{HIGH}$/Input and KRAS$^{HIGH}$/KRAS$^{LOW}$ cell fractions representing negative regulators of KRAS protein abundance. **(B)** ENRICHR-based REACTOME pathway enrichment analysis (*58*) of significant gene candidates identified in the KRAS$^{HIGH}$/Input cell fraction. Most significantly enriched pathways are highlighted.

### Figure S5

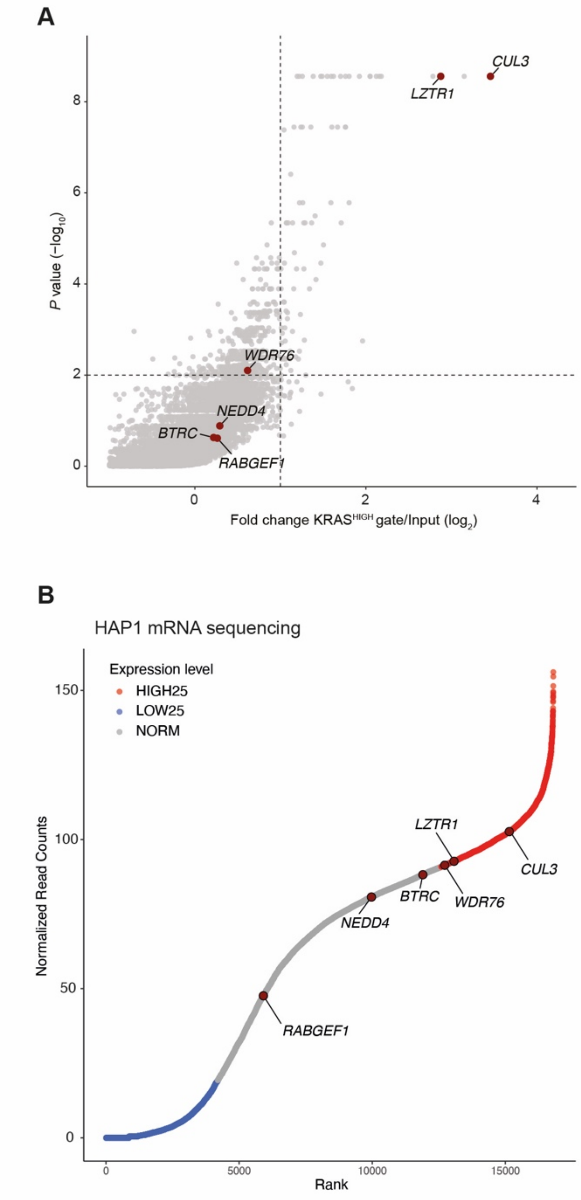


**Figure S5. Functional genetic identification of KRAS abundance-regulating E3 ubiquitin ligases. (A)** Gene-level enrichment of sgRNAs identified in KRAS$^{HIGH}$/Input cell fractions representing negative regulators of KRAS protein abundance. Candidate genes encoding E3 ubiquitin ligases that have been described in the literature to regulate the main RAS GTPase family are highlighted*(8–11,22–25)*. **(B)** Ranked expression of E3 ubiquitin ligases genes as described in **(A)** in HAP1 cells. mRNA expression is derived from published RNA sequencing data(*48*).

**Figure S6**

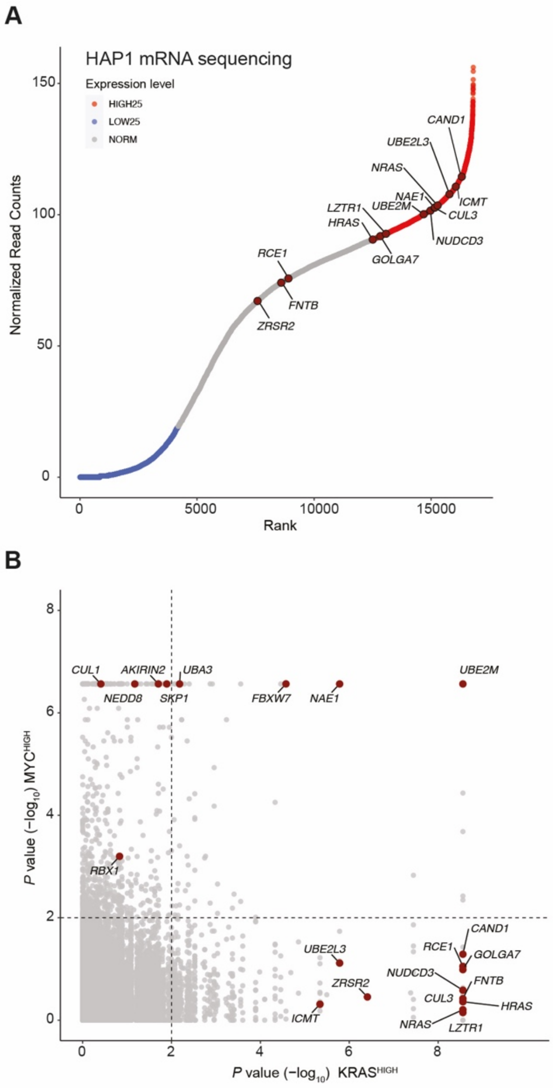


**Figure S6. Functional genetic analysis of proteostatic KRAS regulation. (A)** Ranked expression of identified KRAS protein abundance positive regulator genes in HAP1 cells. mRNA expression is derived from published RNA sequencing data(*48*). **(B)** Comparative analysis of FACS-based CRISPR screens for negative regulators of KRAS and MYC protein abundance. Gene-level enrichment of sgRNAs identified in the $MYC^{HIGH}$ cell fraction in RKO cells is derived from published CRISPR screens(*26*).

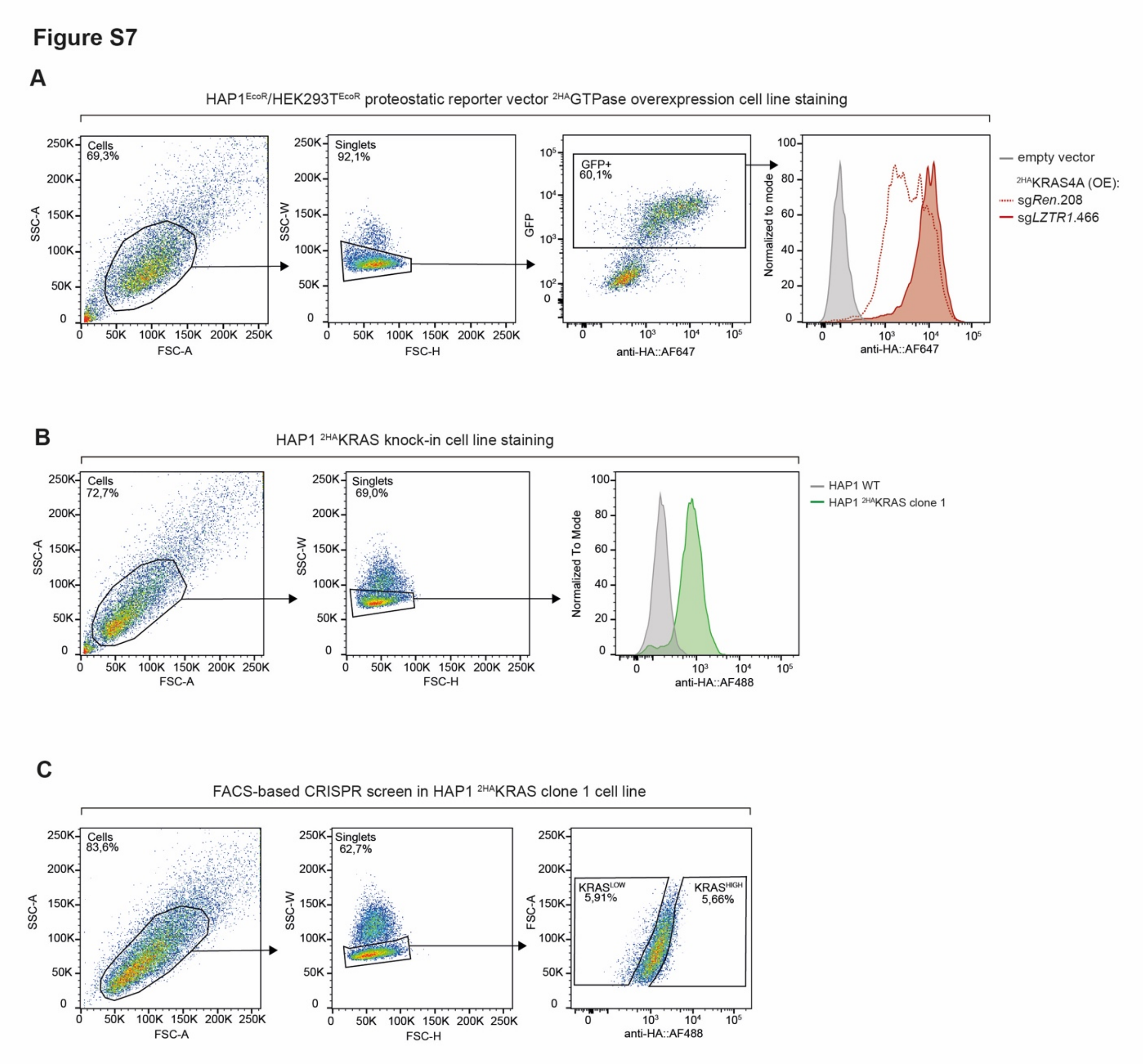


**Figure S7. Gating strategy for flow cytometry analysis and cell sorting.** Representative scatter plots of hierarchical gating strategies. **(A)** HAP1$^{EcoR}$ and HEK293T$^{EcoR}$ proteostatic reporter vector $^{2HA}$GTPase overexpression cell line staining. **(B)** HAP1 $^{2HA}$KRAS knock-in cell line staining. **(C)** FACS-based CRISPR screen in HAP1 $^{2HA}$KRAS clone 1 cell line. Debris and cell doublets were excluded based on forward- (FSC) and sideward-(SSC) scatter.

**Table S1. Primer sequences used for CRISPR screen Illumina library preparation.**

**Table S2. FACS-based CRISPR screen results of KRAS abundance regulation.** MAGeCK-based gene-level enrichment of sgRNAs identified in $KRAS^{HIGH}$/Input, $KRAS^{LOW}$/Input and $KRAS^{HIGH}$/$KRAS^{LOW}$ cell fractions representing negative and positive regulators of KRAS protein abundance. Additionally, MAGeCK-based gene-level enrichment of sgRNAs identified in unsorted input cell fractions compared to initial CRISPR library plasmid DNA (pDNA) has been used to identify essential gene candidates. Result table of ENRICHR-based REACTOME pathway enrichment analysis of significant gene candidates identified in the $KRAS^{HIGH}$/Input cell fraction.

**Supplementary information.** Insert sequences of constructs used for PITCh/MMEJ-based knock-in KRAS cell line generation.

2HA*KRAS*-PITCh-knock-in v1 (KRAS WT):

gcatcgtacgcgtacgtgtttggactgaatataaacttgtggtcaccatggccaagcctttg
tctcaagaagaatccaccctcattgaaagagcaacggctacaatcaacagcatccccatctc
tgaagactacagcgtcgccagcgcagctctctctagcgacggccgcatcttcactggtgtca
atgtatatcattttactggggaccttgtgcagaactcgtggtgctgggcactgctgctgct
gcggcagctggcaacctgacttgtatcgtcgcgatcggaaatgagaacaggggcatcttgag
cccctgcggacggtgccgacaggtgcttctcgatctgcatcctgggatcaaagccatagtga
aggacagtgatggacagccgacggcagttgggattcgtgaattgctgccctctggttatgtg
tgggagggcggatccggcgcaacaaacttctctctgctgaaacaagccggagatgtcgaaga
gaatcctggaccgatgtacccctacgacgtgcccgactacgccggctatccgtatgatgtcc
cggactatgcaggaagcacggagtacaagctggttgtagttggagctggtggcgtagccaaa
cacgtacgcgtacgatgc

2HA*KRAS*-PITCh-knock-in v2 (KRAS WT, G12D and G12C):

gcatcgtacgcgtacgtgtttgggtggtagttggagctggtgggatggccaagcctttgtct
caagaagaatccaccctcattgaaagagcaacggctacaatcaacagcatccccatctctga
agactacagcgtcgccagcgcagctctctctagcgacggccgcatcttcactggtgtcaatg
tatatcattttactggggaccttgtgcagaactcgtggtgctgggcactgctgctgctgcg
gcagctggcaacctgacttgtatcgtcgcgatcggaaatgagaacaggggcatcttgagccc
ctgcggacggtgccgacaggtgcttctcgatctgcatcctgggatcaaagccatagtgaagg
acagtgatggacagccgacggcagttgggattcgtgaattgctgccctctggttatgtgtgg
gagggcggatccggcgcaacaaacttctctctgctgaaacaagccggagatgtcgaagagaa
tcctggaccgatgtacccctacgacgtgcccgactacgccggctatccgtatgatgtcccgg
actatgcaggaagcactgaatataaacttgtggtggtgggcgctggaggcgtaggcaagagt
gccttgaccaaacacgtacgcgtacgatgc

| | |
|---|---|
| 5’ and 3’ PITCh sgRNA target site | NNN |
| 5’ and 3’ 20nt *KRAS* microhomology | NNN |
| Blasticidin resistance sequence | NNN |
| P2A peptide | NNN |
| 2HA tag | NNN |
| KRAS cDNA add-back | NNN |
| WT, G12D, G12C residue | NNN |
